# A Reaction-Diffusion Model for Radiation-Induced Bystander Effects

**DOI:** 10.1101/094375

**Authors:** Oluwole Olobatuyi, Gerda de Vries, Thomas Hillen

**Affiliations:** Centre for Mathematical Biology, Mathematical and Statistical Sciences, University of Alberta, Edmonton, T6G2G1, Canada E-mail: E-mail

**Keywords:** Radiation-induced bystander signal, Cytochrome complex, Bystander effects, Hyper-radiosensitivity, Increased radioresistance, Reaction-diffusion model

## Abstract

We develop and analyze a reaction-diffusion model to investigate the dynamics of the lifespan of a bystander signal emitted when cells are exposed to radiation. Experimental studies by Mothersill and Seymour 1997, using malignant epithelial cell lines, found that an emitted bystander signal can still cause bystander effects in cells even 60h after its emission. Several other experiments have also shown that the signal can persist for months and even years. Also, bystander effects have been hypothesized as one of the factors responsible for the phenomenon of low-dose hyper-radiosensitivity and increased radioresistance (HRS/IRR). Here, we confirm this hypothesis with a mathematical model, which we fit to Joiner’s data on HRS/IRR in a T98G glioma cell line. Furthermore, we use phase plane analysis to understand the full dynamics of the signal’s lifespan. We find that both single and multiple radiation exposure can lead to bystander signals that either persist temporarily or permanently. We also found that, in an heterogeneous environment, the size of the domain exposed to radiation and the number of radiation exposures can determine whether a signal will persist temporarily or permanently. Finally, we use sensitivity analysis to identify those cell parameters that affect the signal’s lifespan and the signal-induced cell death the most.

## 1 Introduction

The radiation-induced bystander effect (RIBE) has fascinated scientists since its first report in 1992 by Nagawasa and Little [32]. They observed that even though less than 1% of irradiated cells were actually traversed by an *α*-particle, about 30% of these cells exhibited radiation effects. At first, RIBE was understood as a secondary effect of radiation on cells exposed to low dose of radiation. These secondary effects of radiation include cell damage, cell death, DNA repair delays and genetic instabilities [38,37]. More experiments [17] have further described RIBE to also include the induction of radiation-like effects in cells that have not been exposed to radiation at all, but are located near an irradiated region. These *by-standing* cells respond to signals — called bystander signals — emitted by irradiated cells and in turn behave as if they have been directly affected by radiation. In fact, it has been shown that these by-standing cells can influence their neighbours and further transport the bystander signal to more distant places. In exceptional cases, the bystander signal has been reported to persist for 31 years in an atomic bomb victim [34] and it was shown in [22] that irradiating a rat’s liver caused a RIBE in its brain and affected the animal’s behavior. These experiments suggest that the RIBE can persist for some time and may also have a non-local behavior.

The nature of the bystander signals is still not fully understood and several mechanisms have been discussed in the literature. For example, ionizing radiation produces free radicals which are, in principle, able to cross cell membranes from cell to cell [5]. However, it is believed that free radicals will quickly react with whatever they encounter and they will not survive very long. Hence free radicals are not able to explain the longevity of the bystander signal. Another explanation considers reactive oxygen species, which can also be transported from cell to cell and react with the DNA[5]. These reactive oxygen species have been identified as bystander signal [1]. Since they behave similar to free radicals, they cannot survive very long as well. Recently however, Cytochrome complex has emerged as a new candidate. Cytochrome complex (cyto-c) plays an important role in oxidative phosphorylation as it transports electrons from the cytochrome b-c1 complex (complex II) to the cytochrome oxidase (complex III) just before the ATP-synthase in complex IV. A functioning glycolysis inside the mitochondria produces energy in the form of ATP, which is essential for the life of the cell. If glycolysis is disrupted, for example due to ionizing radiation, cyto-c might leave the mitochondria, diffuse through the cell, and interact with other proteins, such as DNA[8]. Since cyto-c is small, it can diffuse, or be transported, to neighbouring cells causing damage to those cells. Cyto-c shows many of the characteristics of a bystander signal, making it a good candidate for further analysis.

In this paper we develop a spatially dependent mathematical model for the dynamics of the bystander signal. Since it is not established that cyto-c is the only bystander signal, we will refer to a general bystander signal. However, the model incorporates the bystander dynamics of cyto-c as far as they are known.

After we develop the new bystander model, we fit it to radiation survival data of Joiner et. al. [18]. Joiner et. al. [18], and many others, found that the cell survival is lower than expected for low-dose radiation exposure. This suggests potential risks associated with low-dose radiation exposure, which may have implications for carcinogenesis and radiation protection. In fact, low-dose radiation exposure has been found to be carcinogenic in some cases [25,16].

The longevity of the bystander signal in tissues has surprised many scientists. Our model allows for a full understanding of the life time of the bystander signal, based on a positive feedback loop. Cells are damaged by radiation, this releases cyto-c into the environment, which damages other cells, which releases more cyto-c into the environment. This feedback will not accumulate cyto-c, but it explains the long persistence of enlarged cyto-c levels.

The model lets us quantify how much tissue damage is related to direct radiation damage versus indirect bystander damage. Not surprising, we find that for low doses the bystander effect is largest, whereas for larger radiation doses, the direct cell kill dominates. These conclusions have also been found in Powathil et. al. [35] using a computational hybrid model.

The rest of this section will further describe some of the important terminology in this paper.

### 1.1 Hyper-radiosensitivity (HRS) and Increased Radio-resistance (IRR)

If ionizing radiation is applied to cells, we consider a cell *clonogenically* dead if it is unable to form cell colonies, i.e., it can no longer produce more than 50 offspring. The (clonogenic) surviving fraction (SF) of cells exposed to radiation is defined as the fraction of the irradiated cells that is capable of forming colonies after radiation exposure. A common model for SF of cells is the linear quadratic (LQ) model

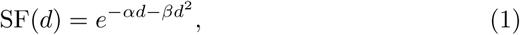

where *d* is the radiation dose (measured in Gy), *α* is the rate at which single radiation tracks produce lethal lesion, and *β* is the rate at which binary mis-repair of pairs of double strand break (DSB) from different radiation tracks lead to lethal lesions [7].

In recent experiments [23,28,43,45,46], it was shown that cell survival deviates from the LQ model prediction for low-dose radiation in the range of 0.1–1 Gy. We illustrate this difference in Figure 1. The measured surviving fraction for low-dose radiation lies significantly below the LQ curve. This phenomenon has been termed low-dose hyper-radiosensitivity (HRS) [18]. The HRS is followed by a period of relative radio-resistance (per unit of dose) of cell kill over the dose range of ~ 0.5 — 1Gy. This later phenomenon is referred to as increased radio-resistance (IRR).

**Fig. 1:**
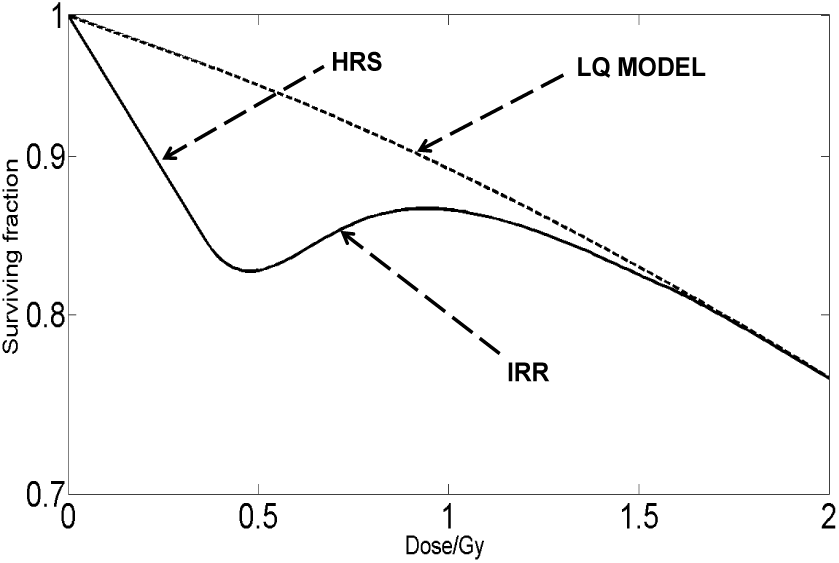
Surviving fraction of cells as a function of dose according to linear quadratic model (1) (dashed line) and with the HRS/IRR phenomenon (solid line)

We will see that our bystander model can describe the observed hyper-radiosensitivity (HRS) and increased radio-resistance (IRR) effects, making the bystander effect a possible explanation of these phenomena.

### 1.2 Bystander Effects

As previously noted, *bystander effects* are secondary effects of radiation either in cells exposed to low dose of radiation or in cells located near an irradiated region [37,38]. Although most bystander effects are observed in direct proximity of the irradiated region, bystander effects have also been reported quite far from the irradiated region [36,22]. Here we will focus on the immediate spatial vicinity of an irradiated region.

The *bystander effects* can be roughly classified into [38,37]:

1. *Bystander signal-induced cell death* Bystander signal-induced cell death occurs when the bystander signal interacts with the DNA of a cell to reduce its proliferation capacity to the extent that it can no longer produce more than 50 offspring. Since bystander signal-induced death may also be other forms of death besides clonogenic death, we will also account for other type of death like apoptosis, necrosis and so on.
2. *Bystander signal-induced cell damage* Bystander signal-induced cell damage occurs whenever the bystander signal interacts with the cell’s DNA to damage its proliferation capacity but the cell can still produce more than 50 offspring.
3. *Cell repair delay^1^* This biological effect occurs when the repair mechanism of damaged DNA of an irradiated cell is interrupted, delayed or completely hindered by the bystander signals [39].
4. *Genetic instability* The interaction of the bystander signal with the DNA can also lead to DNA disrepair or mutations. We will not include genetic instabilities in our mathematical model, since these do not contribute to the dynamics of the bystander signal. Mutations might create cancers, or promote cancer development, but this is beyond the scope of this paper.

As mentioned above, several molecules are discussed as bystander signals, and cyto-c is currently the most convincing candidate. Since this signal can diffuse from cell to cell it is possible that irradiated cells are also affected by the bystander signal. The release of cyto-c after radiation depends on the tumor suppressor gene p53. p53 initiates DNA repair but, at the same time, p53 initiates apoptosis when damage is irreparable. This apoptotic pathway basically involves destruction of the mitochondria, releasing cyto-c into the cytoplasm [41,12,11,10]. When cyto-c is released from a damaged cell, it is transmitted either via gap junctions [3] to neighboring cells or it diffuses to neighboring cells [33]. The cyto-c protein can initiate double strand breaks (DSBs) in neighboring cells [20]. These DSBs in neighboring cells also activate p53, which eventually can lead to further release of cyto-c. In this way the cycle continues as illustrated in Figure 2 and we can obtain a rather long-lived signal cascade.

**Fig. 2:**
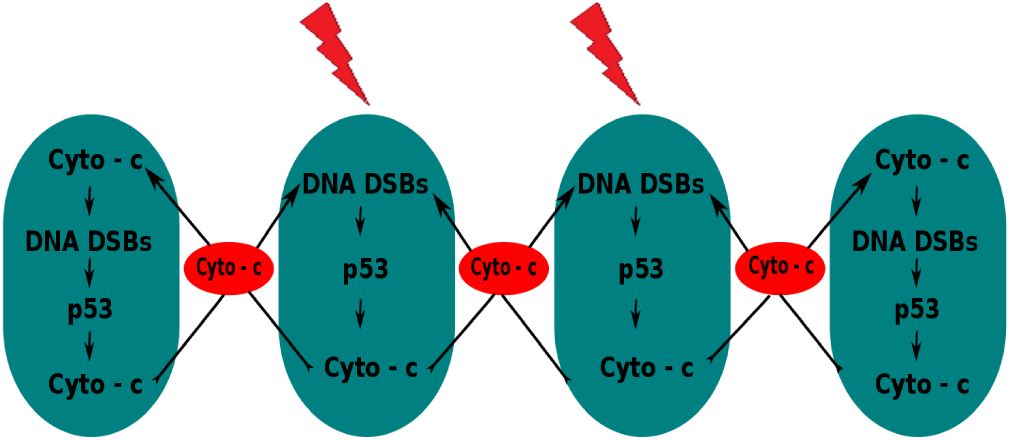
p53-induced coupling between irradiated cells and between irradiated and un-irradiated cells. The green ellipsoidal structures represent cells while the lightning symbols indicate radiation.

### 1.3 Bystander Signal’s Lifespan

Bystander effects have been found in various *in vivo* and *in vitro* experiments to outlive the direct radiation effects. Pant and Kamada [34] reported the presence of bystander signal in the blood plasma of heavily exposed atomic bomb survivors 31 years after exposure. They found the presence of these signals when blood cells of normal individuals exhibited bystander effects when they were cultured with the blood cells of bomb survivors. Similarly, Goyanes-Villaescusa [15] reported the presence of bystander signals in the children born to mothers exposed to radiation several months before conception. These observations have led to an open question regarding “How long does a bystander signal live after its emission?”[6]. Our mathematical model will allow us to address this question in detail. We find that a saddle point in the phase plane of the bystander model is the organizing centre for the longevity of the bystander signal. Orbits pass by a saddle point, hence leading to a long transient.

### 1.4 Previous Models of Bystander Effects

The idea of modelling radiation-induced bystander effects using diffusion-based mechanism began with Khvostunov et. al. [19,33]. These early models use a quantitative approach that assumes that bystander signal is a protein-like molecule spreading via diffusion. The bystander effects considered are cell death and mutation induction. They assume that bystander effects only occurs in un-irradiated neighboring cells and the bystander response to this signal is assumed to be a binary “on/off” mechanism. They found that their bystander model can explain the experimental data for cell survival and induced oncogenic transformation frequencies. These data also confirm the assumption of the protein-like nature for the bystander signal.

In 2013, McMahon et. al. [29] first modeled the evolution of bystander signal with a partial differential equation (PDE) while a computational model was used to describe bystander cell’s responses. Their model also assumes a binary cell’s response to bystander signal but incorporates the occurrence of bystander effects in the irradiated cells as well. Their model only incorporated bystander signal-induced damage, which is assumed to result into cell death, cell-cycle arrest or further cell damage. This model suggests that bystander effects play a significant role in determining cellular survival, even in directly irradiated populations.

Most influential to our work is a recent work (2016) by Powathil et. al. [35]. They developed a hybrid multi-scale mathematical and computational model to study multiple radiation-induced bystander effects (bystander signal-induced damage and death, and cell repair delay) on growing tumor within host normal tissue. In their model, the evolution of the bystander signal was described by a PDE, while a stochastic process was used to describe cellular evolution and responses to the signals. They also assumed that both irradiated and un-irradiated cells can respond to the bystander signals. They assumed that bystander signals are emitted by radiation-induced dead and damaged cells. Their model shows that bystander responses play a major role in mediating radiation damage to cells at low-doses of radiation, doing more damage than that due to direct radiation. The survival curves derived from this shows an area of hyper-radio-sensitivity at low-doses that are not obtained using a traditional radio-biological model.

In our model, we use PDE to describe both the dynamics of cells and the signals. In general, we extend most of the assumptions of Powathil et. al. in [35] but in a continuous setting. With the recent discovery of cyto-c as a candidate for the bystander signal, our model is able to incorporate more biologically justifiable assumptions. In particular, we assume that damaged cells do not proliferate since they are expected to be under cell cycle arrest. This assumption was not incorporated in any of the previous models. We also assume that cells which are damaged via bystander signals can also emit bystander signal. This is because DNA damage, irrespective of the cause, can trigger the activity of p53 which eventually leads to the emission of cyto-c as seen in Figure 2. The model also assumes that damaged cells emit bystander signals as long as they are not repaired or dead. It is more biological to model cellular response to bystander signal with a continuous functional rather than a discontinuous binary “on/off” mechanism. Finally, both irradiated and unirradiated cells can exhibit the bystander effects.

Our model agrees with previous results on bystander effects both experimental and theoretical. In particular, our bystander model shows that bystander effects are predominant at low-doses of radiation and it is a major contributor to the phenomenon of hyper-radiosensitivity. Furthermore, we use our model to determine cell parameters that affect this phenomenon the most. Also, we use our model to extensively study how long an emitted signal lives and to show that, although bystander signal is produced at every radiation exposure, it does not increase with increase in radiation exposure. In fact, we found that the concentration converge to a steady state after a couple of exposures.

### 1.5 Outline of the paper

In Section 2, we formulate and develop the mathematical model, explain the relevant parameters and parameter functions, and present a nondimensionalization of the model. We also fit the model to the radio-sensitivity data of Joiner [18] and estimate the model parameter values as well as their 95% confidence intervals. In Section 3, we numerically explore the qualitative behavior of the bystander signal profile when cells undergo both single and multiple radiation exposures, respectively, in both homogeneous and heterogeneous domains. The numerical exploration in Section 3 shows that the bystander signal can live quite long, hence in Section 4 we analyze the bystander signal lifespan. In the analyses on the signal’s lifespan in Section 4, we perform a phase-plane analysis to identify a saddle point as the organizing centre for the bystander signal dynamics. We also perform a sensitivity analysis to see which parameters have the most impact on the lifespan of the signal. After having gained a good understanding of the bystander dynamics, we analyze the relative significance of the bystander effect on cell survival in Section 5. We show that this effect is significant for small radiation doses (0.1 – 1 Gy), but it is almost irrelevant at larger doses. We finish the paper with a discussion in Section 6.

## 2 The Bystander Model

### 2.1 Model Development

The mathematical model of this paper describes the biological interactions between healthy cells, damaged cells, radiation energy, and bystander signals. We base the model on a typical bystander - *in vitro* assay [17], where cells are cultured on a petri-dish in a monolayer and radiation is applied to a certain partial area of the dish.

A schematic of the model, illustrating the relationship between the components, is given in Figure 3. The density of healthy cells is denoted by *u*, the density of damaged cells by *v* and the concentration of bystander signal by *w*. In Figure 3, we indicate that a cell exposed to ionizing radiation can either be damaged, or killed, or be completely unharmed and reproduce [24]. After radiation exposure, the fraction of the cells that is damaged by radiation enters the damaged cell compartment *v*. As suggested by the biology of cyto-c, we assume that radiation-induced damaged cells emit bystander signal as long as they live, while dead cells release bystander signal into the environment once at the time of their death.

**Fig. 3:**
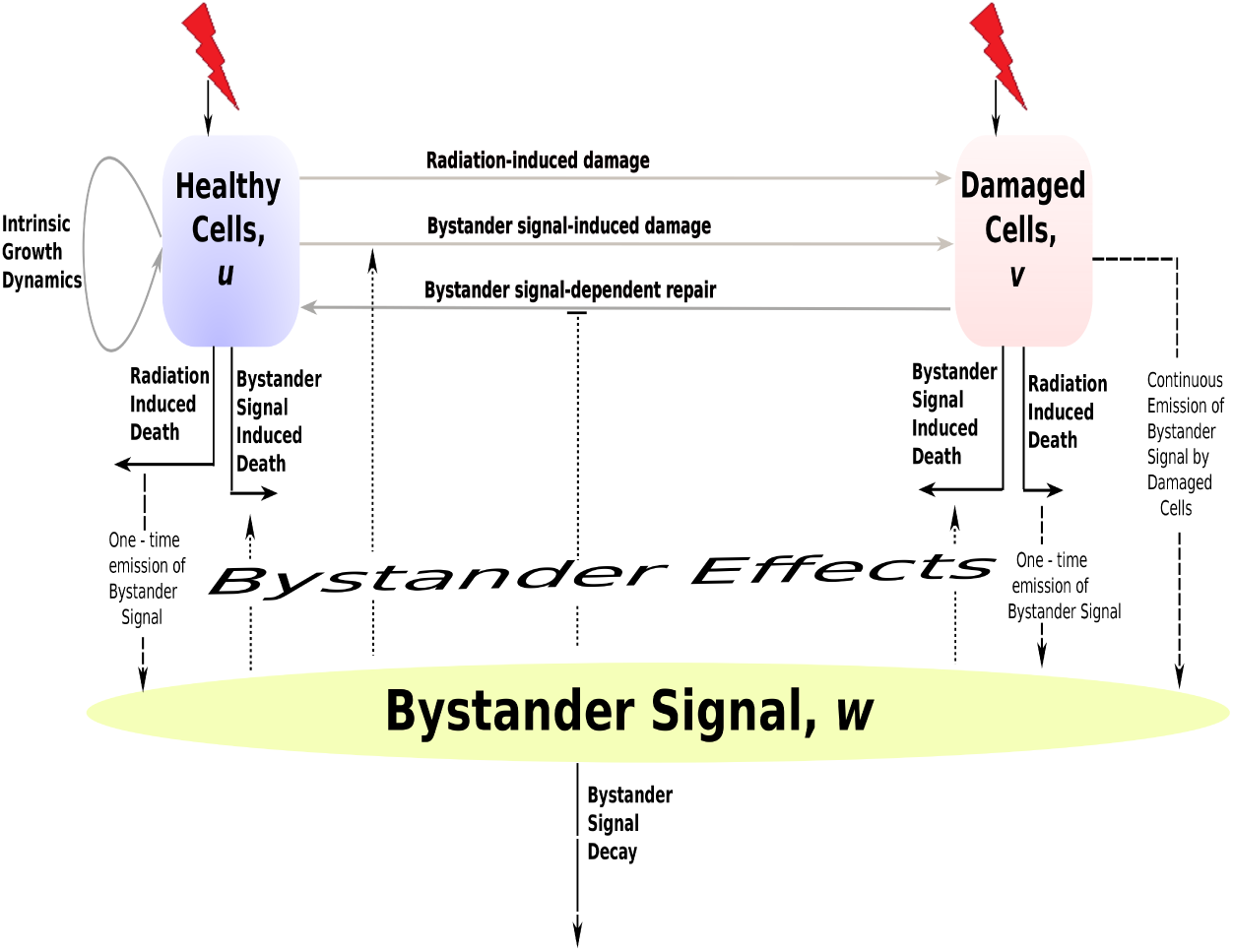
Schematics of the bystander signal model. The dotted lines represent the bystander effects, the dashed line represents the bystander signal emission, the grey solid line represent cellular interactions, and the black solid lines represent cellular responses to both radiations and bystander signals. The lightning symbols indicate radiation.

Whenever a cell’s DNA is damaged, several mechanisms are triggered to repair the damage [39,2]. Thus, damaged cells have the potential to fully recover and become healthy cells again. The emitted bystander signals, *w*, provide negative feedback to both the healthy and the damaged cells by causing cell death, cell damage, and DNA repair delay. If a damaged cell is further damaged, then we keep it in the class of damaged cells until it either dies or is repaired. Finally, the bystander signal is produced by dying and damaged cells, it diffuses through the petri dish and it can decay or be cleared out.

To formulate the model mathematically, we denote the number of healthy cells by *u*(*x*,*t*), the number of damaged cells by *v*(*x*,*t*), and the concentration of the bystander signal by *w*(*x*, *t*). Here *x* ∊ *Ω* denotes the spatial coordinate in a smooth bounded domain *Ω* ⊂ ***R***^*n*^ and *t* ≥ 0 denotes time. The model that describes the mutual interactions between *u*, *v*, and *w* based on the schematics described in Figure 3 is given by:

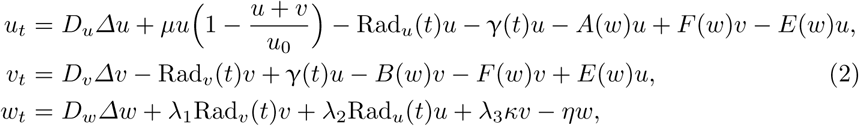

with appropriate boundary conditions depending on the biological situation.For instance, Neumann boundary conditions will be suitable for the petri dish experiment. Details are as follow:

1. In this model, all components are subject to spatial diffusion, expressed through the Laplacians Δ*u*, Δ*v*, and Δ*w*. In case of healthy normal tissue, the diffusion coefficients *D*_*u*_ and *D_v_* might be zero, but if applied to tumor tissue, then *D_u_* and *D_v_* will be nonzero due to local spread of the tumor. Since the signal *w* can diffuse everywhere, we assume a nonzero diffusion coefficient *D_w_* > 0.
2. Healthy cells, *u*, grow according to a logistic law with carrying capacity, *u*_0_, and growth rate *μ*.
3. Biological cells respond to radiation exposure in two ways namely: radiation-induced death at rate Rad*_i_*(*t*) (for *i* = *u*, *v*) and radiation-induced damage at rate *γ*(*t*). We model the radiation-induced death rate by the radiation hazard function for fractionated radiation [13]

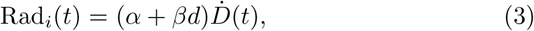

where *d* is the dose per fraction, *Ḋ*(*t*) denotes the dose rate (i.e. a piecewise constant jump function), *α* is the rate at which single radiation tracks produce lethal lesion, and *β* is the rate at which binary misrepair of pairs of double strand break (DSB) from different radiation tracks lead to lethal lesions [7].
4. To model the radiation-induced damage rate γ, we consider the classical Lethal-Potentially Lethal Model (LPL) of Curtis [9] for radiation damage. As shown in Figure 4 (solid curve), the damage rate in the LPL model has a characteristic shape of a steep increase, a single global maximum and a slow decay to zero. We describe this behaviour by the function shown in Figure 4 (dotted curve), which is given by

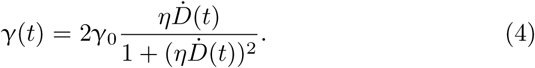 Here *γ*_0_ is the maximum damage rate and *η* = 1/*Ḋ*_max_ denotes the reciprocal of the dose rate *Ḋ*_max_ at which the radiation damage is maximal.
5. The bystander effects in system (2) are described by the functions *A*(*w*), *B*(*w*), *E*(*w*), and *F*(*w*). The rates *A*(*w*) and *B*(*w*) denote the bystander signal-induced death rates for healthy and damaged cells, respectively. *E*(*w*) denotes the bystander signal-induced damage rate and *F*(*w*) denotes the bystander signal-dependent damage repair rate. Mothersill and Seymour suggested in [42, 40] that responses to bystander signals are threshold phenomena. The signal induces a response only when the signal concentration exceeds a lower threshold, and it reaches a maximum response at an upper threshold. All the bystander effects in this paper will be modeled with two threshold parameters (see Figure 5) using a generic tanh profile:

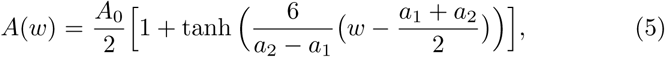

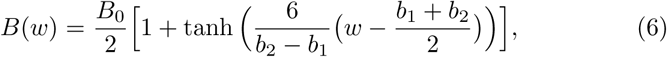

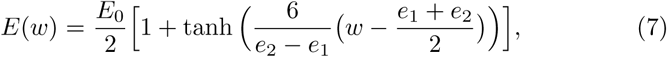

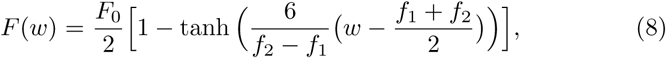 These models contain threshold values *a*_1_ < *a*_2_, *b*_1_ < *b*_2_, *e*_1_ < *e*_2_, and *f*_1_ < *f*_2_ respectively. The repair rate, *F*(*w*), of damaged cells is a decaying function of *w*, indicating reduced repair capabilities if *w* exceeds the lower threshold *f_1_*. The above four rate functions are plotted in Figure 5.

**Fig. 4:**
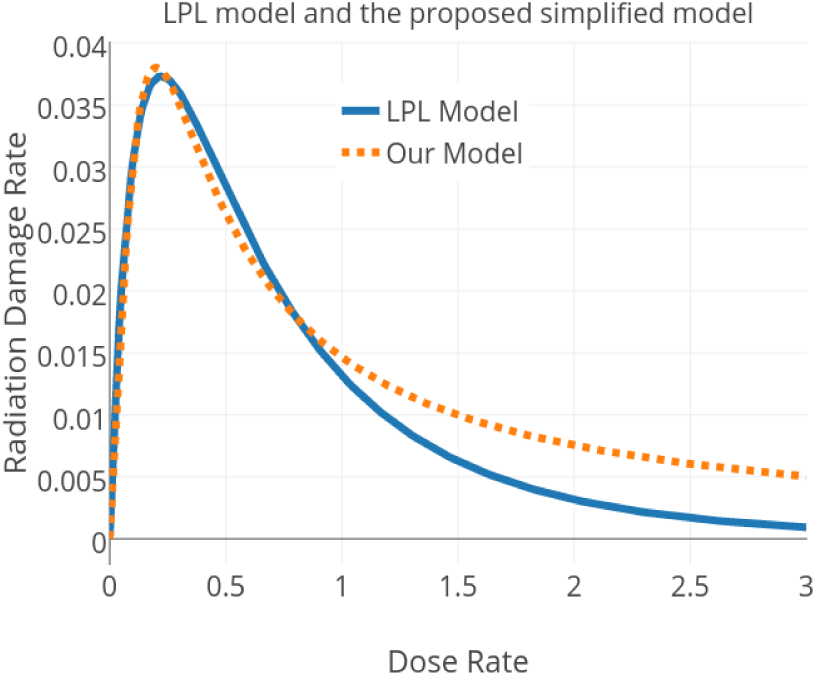
The profile of radiation-induced damage rate, *γ*(*t*), according to both the LPL model and our proposed simplified model. The solid line corresponds to the LPL model [9] while the dotted line corresponds to our proposed simplified model for radiation damage rate at *γ*_0_ =0.038 and *d*_max_ = 0.2 Gy.

**Fig. 5:**
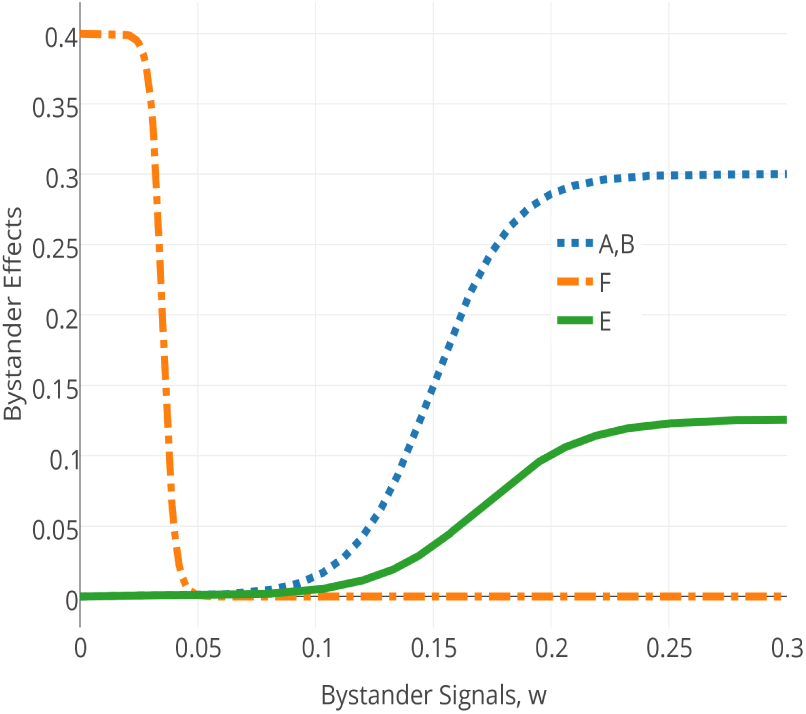
Bystander Effects, *A*, *B*, *E*, and *F*. The parameter values are *a*± = *b*_1_ = 0.05, *a*_2_ = *b*2 = 0.25 & *A*_0_ = *B*_0_ = 0.3 for *A*(*w*) and *B* (*w*), *f*_1_ = 0.02, *f_2_* = 0.05 & *F*_0_ = 0.4 for *F* (*w*), *e*_1_ = 0.04, *e*_2_ = 0.3 & *E*_0_ = 0.1333 for *E* (*w*).

### 2.2 Rescaling

The bystander signal model (2) contains many parameters. The parameters whose values are found in the literature are listed in Table 1. The remaining parameters will be estimated from Joiner’s data set in the next subsection. However, before we attempt to estimate the parameters, we will rescale the model to carrying capacity *u*_0_ = 1 and unit decay rate of the bystander signal.

**Table 1:**
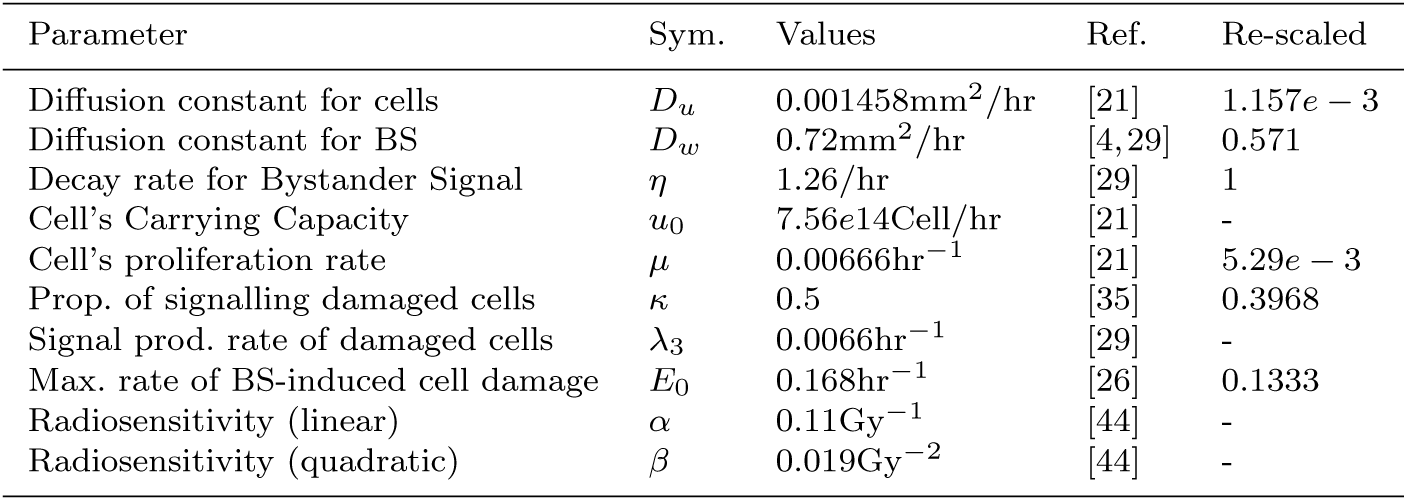
Parameter values available in the literature and their references.

We apply the following transformations

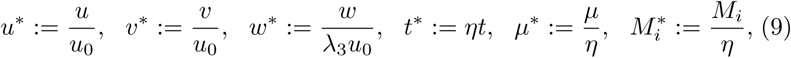

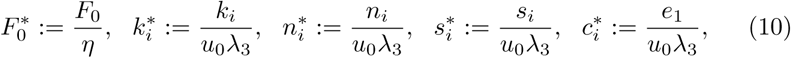

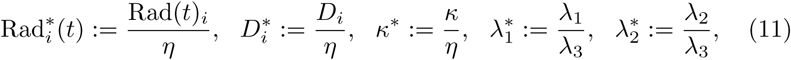

to model (2) and arrive at the following dimensionless model after dropping the asterisk (*)

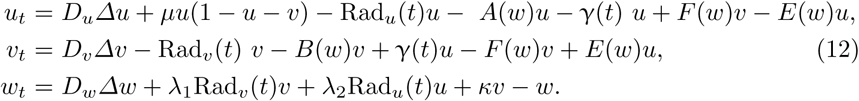

### 2.3 Parameterizations and Data Fitting

The parameters that are unknown (see Table 2) are mostly parameters such as the bystander effect thresholds which may be difficult to measure directly. Thus, there is the need to accurately estimate their values and a confidence interval for each of them, which will form the parameter space for our model.

The data of Joiner et. al. [18] describe the survival of asynchronous T98G human glioma cells irradiated with 240kVp X-rays. These cells were irradiated with single doses of X-ray between 0.05 and 6 Gy at a dose rate of 0.2–0.5 Gy per min. The surviving fraction of cells following exposure to single doses was measured using a Cell Sorter. Each data point in Figure 6 represents between 10–12 measurements and is denoted as mean ± the standard deviation. Cell survival was described in terms of their ability to form a colony (i.e. reproduce at least 50 offspring after radiation exposure) and cells which are unable to form a colony do not survive.

**Fig. 6:**
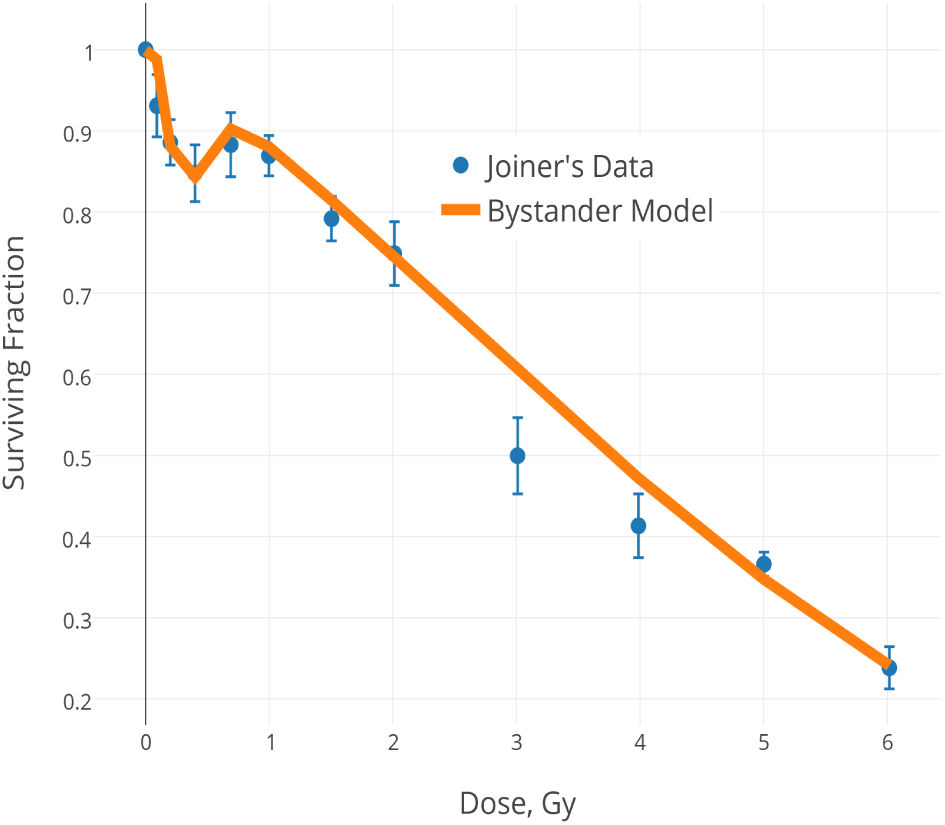
Data fitting. The dots with error bars are the data from Joiner et. al. [18] while the solid curve is the fit from the bystander signal model.

We employ an implementation of the Goodman and Weare Affine invariant ensemble Markov Chain Monte Carlo (MCMC) sampler [14] to fit the model to the dataset in [18]. In particular, we fit the surviving fraction, SF(*t*), of irradiated cells at time *t* = 6h computed from the bystander signal model (12) to the surviving fraction data [18]. The affine invariance property of this routine enables a much faster convergence even for badly scaled problems. The implementation takes a log-likelihood function of the experimental data and a log-prior of each parameter as input. We assume an exponential distribution for these data. We also assume a uniform distribution for the prior of each parameter over the prescribed intervals of biologically relevant values. The implementation of this MCMC sampler on the bystander signal model results in the fit shown in Figure 6. The values of the parameters and their respective 95% confidence intervals are given in Table 2 below.

**Table 2:**
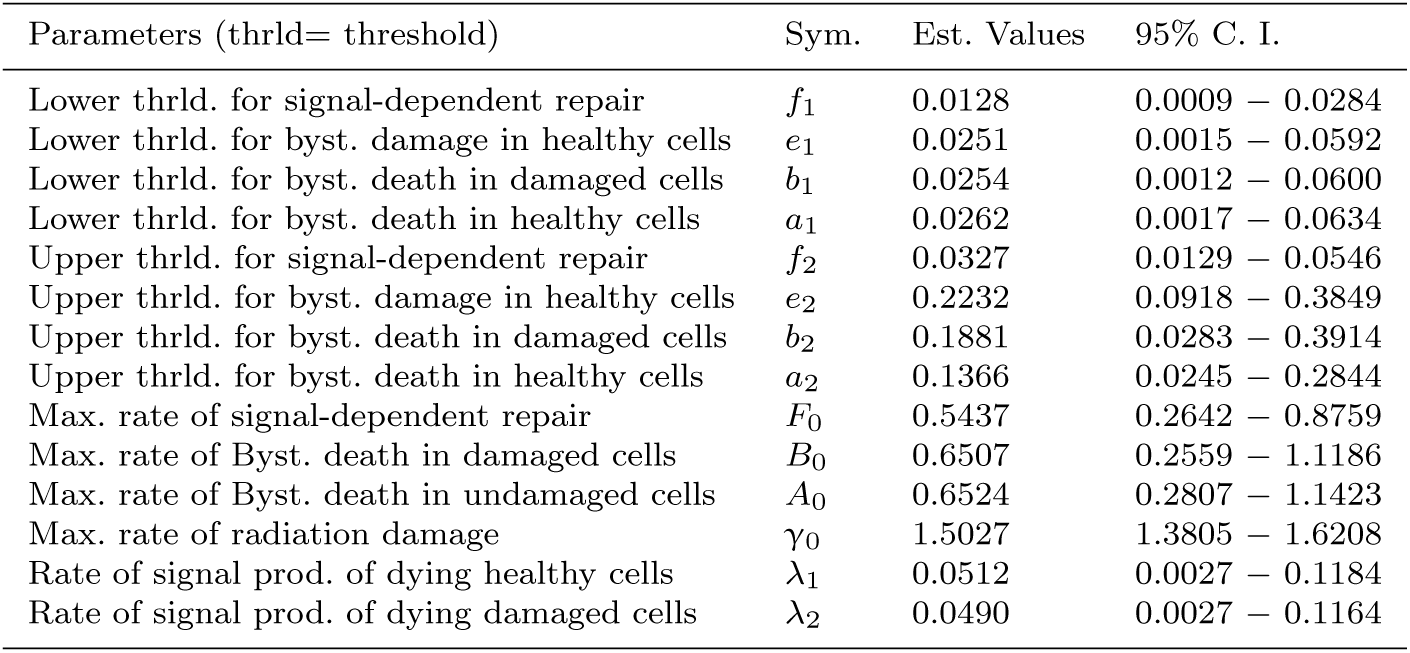
Parameters and their values estimated using an MCMC sampler.

We need to reiterate that the parameters estimated are the dimensionless parameters. For example, we will interpret the estimated threshold values in unit of bystander signal per unit of cell’s carrying capacity per unit of rate of bystander signal production. The 95% confidence intervals of these parameter values will allow for further study of the bystander signal’s dynamics via mathematical techniques like bifurcation and sensitivity analyses.

This parameter estimation gives some very useful insight into the parameter relationships between the healthy and damaged cells. For example, the estimation suggests that the bystander signal-induced death rate and the rate of one-time bystander signal emission by radiation-induced dead cells are the same in both the healthy and damaged cells. Thus, we will henceforth assume that *A* = *B* and λ_1_ = λ_2_. This estimation also suggests that the dynamics of the bystander signal is faster than the cellular dynamics. This fact will become important in Section 4 as we reduce the system of three PDEs to a system of two ODEs for phase plane analysis.

As noted earlier, this system of three PDEs was motivated by the stochastic spatial model of Powathil et. al. [35]. Thus, in the next section, we will numerically explore this system of PDEs, although much of the qualitative behavior can be captured by the corresponding system of ODEs, as we will show in Chapter 4. The standard parameter set used for all the simulations in this paper consists of the parameter values found in the literature (which are highlighted in Table 1) and the default parameter set in Table 3 is a subset of the dimensionless parameter space (from Table 2).

**Table 3:** Default parameter set for the simulation in this paper.

| $a_1$ | $a_2$ | $A_0$ | $e_1$ | $e_2$ | $f_1$ | $f_2$ | $F_0$ | $\lambda_1$ | $\gamma_0$ |
| --- | --- | --- | --- | --- | --- | --- | --- | --- | --- |
| 0.05 | 0.25 | 0.3 | 0.04 | 0.3 | 0.02 | 0.05 | 0.4 | 0.016 | 1.5 |

## 3 Numerical Exploration of the Model

For the numerical exploration of the model (12), we choose parameter values from Tables 1 and 3. We use Matlab to solve system (12) on an interval [0,1] with spatial step *dx* = 0.05 and temporal step *dt* = 1/60 (corresponding to 1 minute). We first consider a single radiation exposure with dose, *d*, given at the beginning of the simulation, which we incorporate by the dose rate

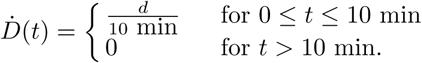

The dose rate is incorporated into both the hazard function (3) and the radiation-induced damage rate (4). Since the bystander effects have been found to be more pronounced at low dose of radiation, we will only irradiate at dose *d* = 0.2 Gy.

We will study the profile of a bystander signal in both homogeneous and heterogeneous domains using the model (12). A domain is termed homogeneous if every part of the domain is *equally* exposed to radiation, otherwise, it is termed heterogeneous. For simplicity, we will restrict our numerical simulations to a one-dimensional domain. We will assume the initial condition (1,0,0), representing cells that are fully grown to its full carrying capacity but have never been previously exposed to radiation. Since we want to study the cellular response at low dose in a petri dish setting similar to [17], it is natural to assume a Neumann boundary condition for all the components. Although the boundary effects on the dynamics of bystander signal are also interesting, we will not consider them in this paper.

We are also interested in studying the behavior of the system under multiple radiation exposures with each exposure corresponding to a daily dose of 0.2Gy at 0.02Gy/min dose rate. Each daily radiation exposure will be referred to as a *fraction.*

### 3.1 Bystander Signal Profile in a Homogeneous Domain

The monolayer cells are exposed uniformly to radiation similar to the setup in [17]. Immediately, we observe a rapid production of the bystander signal reaching a maximum shortly after the exposure as illustrated by Fig. 7a. Although the signal concentration slowly declines, it seems to converge to some nonzero concentration at *w* ≃ 0.0653. This value is higher than the lower thresholds for all the bystander effects. This suggests that even in the long run, the bystander effects can persist. On the other hand, if the maximum rate of bystander signal-induced death, *A*_0_, in cells is increased to 0.7 as seen in Fig. 7b, we observe that the signal concentration is washed out at some later time after radiation exposure. This observation about the dynamics of the signal’s lifespan raises the question of ‘How long can bystander effects persist in cells after radiation exposure and what are the parameters that affect the dynamics?’ We will fully answer this question in Section 4 using phase plane and sensitivity analyses.

**Fig. 7:**
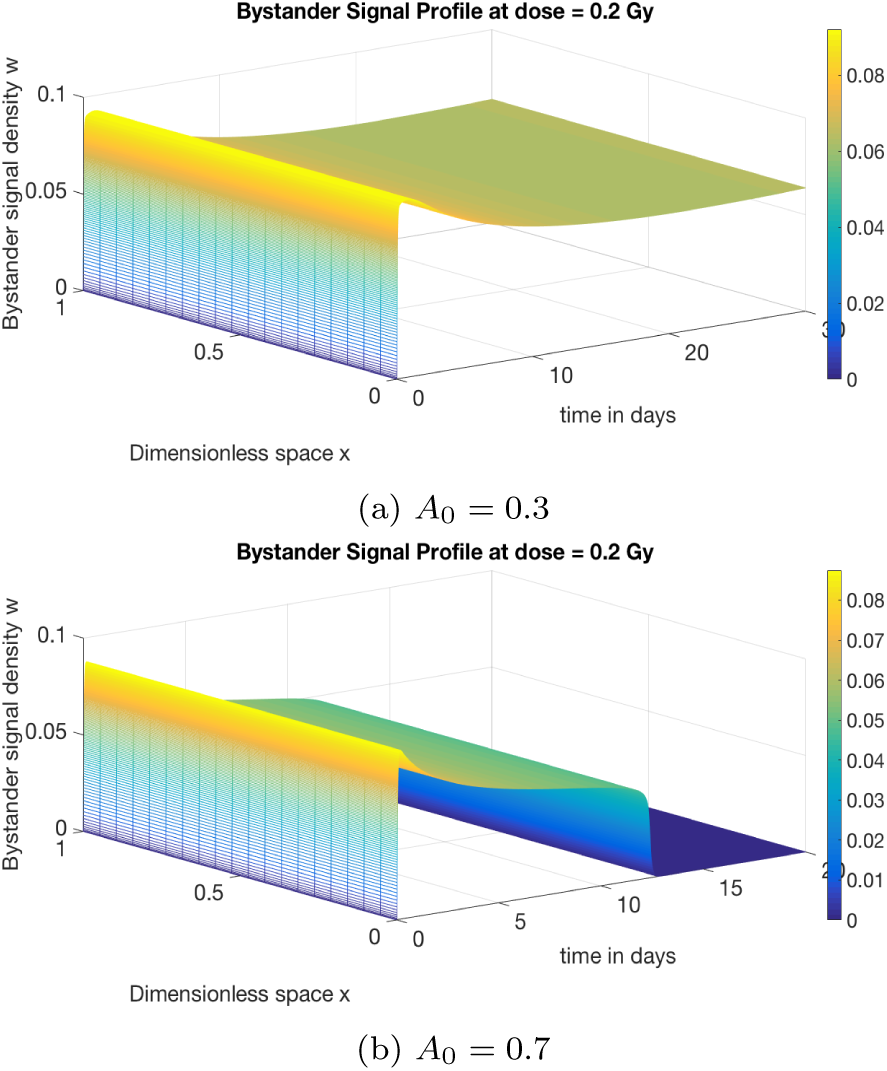
Bystander signal profiles with both single radiation exposure at different values of *A*_0_.

The profile of the emitted bystander signal is quite different under multiple radiation exposures. At each fraction, there is an increase in the signal’s concentration with the maximum concentration occurring immediately after the second fraction as seen in Fig. 8. In fact, further exposure to radiation after about 7d does not significantly increase the concentration of the signal as seen in Fig. 8c.

**Fig. 8:**
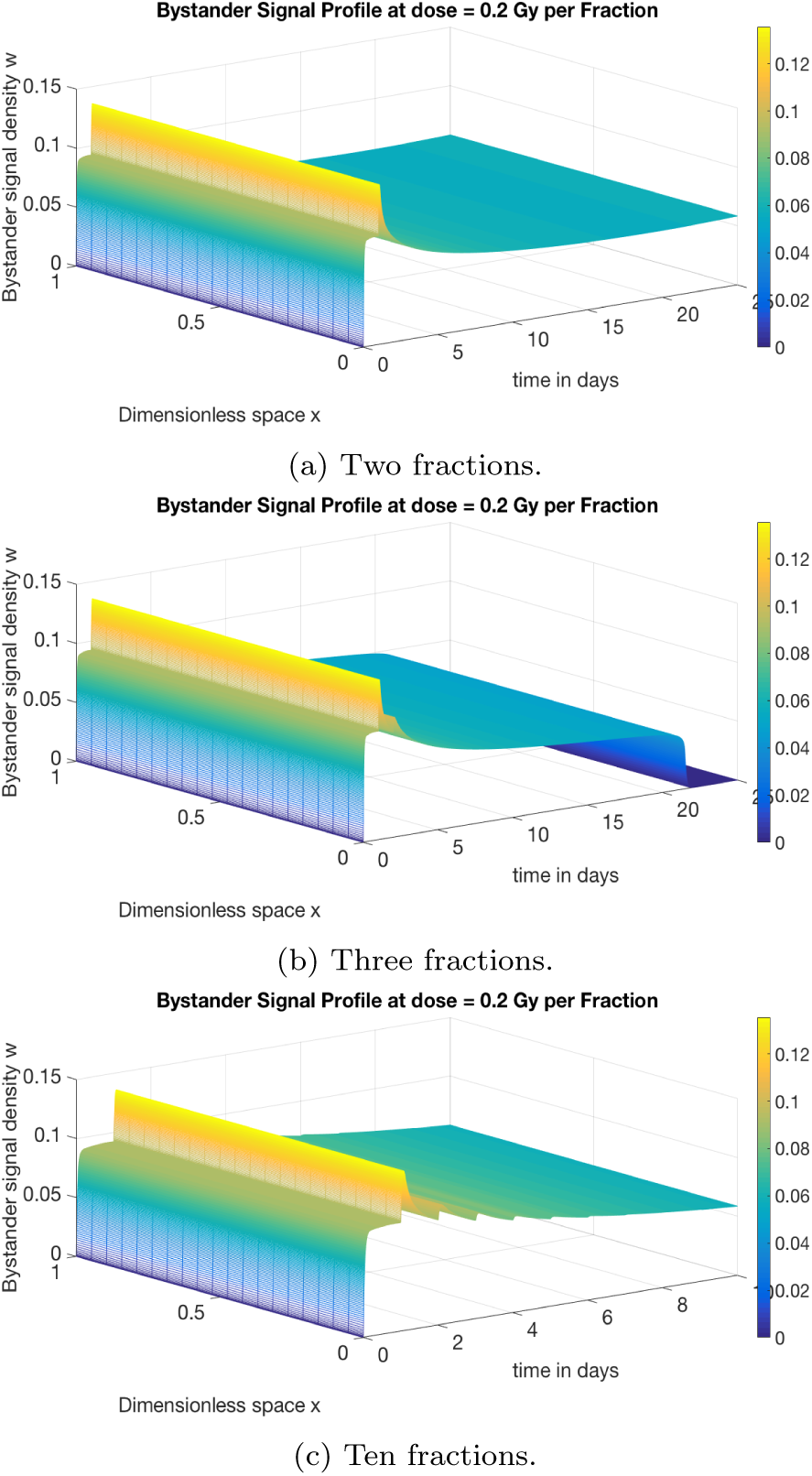
Bystander signal profiles with multiple radiation exposures at *A*_0_ = 0.3.

We noted in Fig. 7a that the long-term dynamics of the signal converge to a nonzero concentration at *A*_0_ = 0.3. However, at the same parameter value, we observe that after three fractions of radiation exposures, the signal converges to zero. Whereas, the nonzero convergence at *A*_0_ = 0.3, is preserved with two fractions as seen in Figure 8a. The number of fractions of radiation exposure appears to affect the long-term dynamics of the signal. We will try to understand this effect on signal dynamics using phase plane analysis in the following section.

### 3.2 Bystander Signal Profile in a Heterogeneous Domain

It is also interesting to study the dynamics of the emitted signal when the cells are not uniformly radiated. In this subsection, all simulations will be done at *A*_0_ = 0.3 unless otherwise stated. Depending on the percentage of the domain exposed to radiation, we observe different dynamics. For instance, when we irradiate ≤ 90% of the domain, even at *A*_0_ = 0.3, the signal converges to zero. Figure 9a corresponds to the system when 90% of the domain is exposed to radiation. Since the same behavior is seen when less < 90% of the domain is irradiated we will not include the simulations. On the other hand, we observe that the bystander signal converges to a nonzero value when ≥ 91% of the domain is exposed to radiation as seen in Figure 9b. This spatially-driven change in the signal’s dynamics is interesting and a rigorous mathematical analysis of this phenomenon is required.

**Fig. 9:**
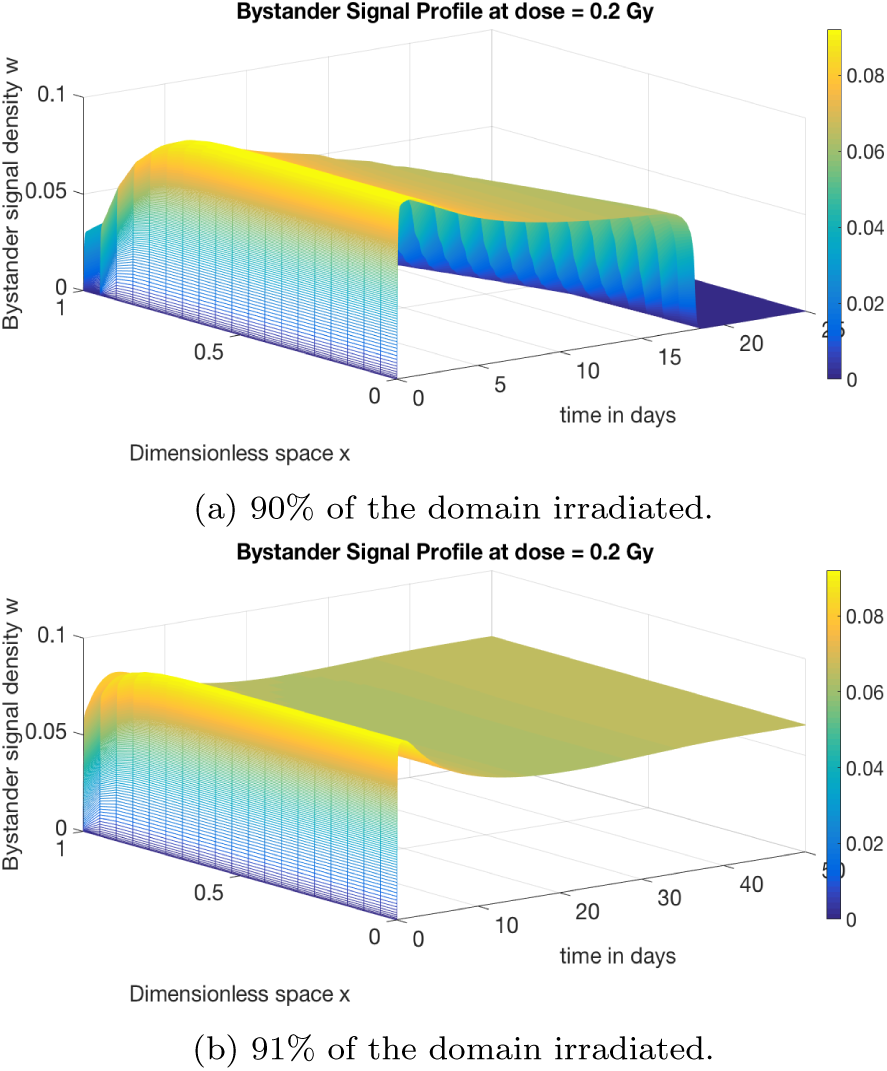
Bystander profiles in a single fraction on a heterogeneous domain at *A*_0_ = 0.3.

The dynamics of bystander signal with multiple radiation exposures in a heterogeneous domain is very interesting due to its application to radiotherapy. For example, tumor undergoing fractionated radiation treatment is usually surrounded by normal cells. We observe that the bystander signal dynamics changes significantly depending on the number of fractional exposures and the percentage of the domain exposed to radiation.

In Figures 10a and 10b we observe the effect of space on the signal dynamics. The signal converges to zero when 85% of the domain is exposed to radiation while the signal converges to a nonzero value when about 86% of the domain is irradiated. We observe this phenomenon as well for a single radiation exposure but with different size of irradiated domain.

**Fig. 10:**
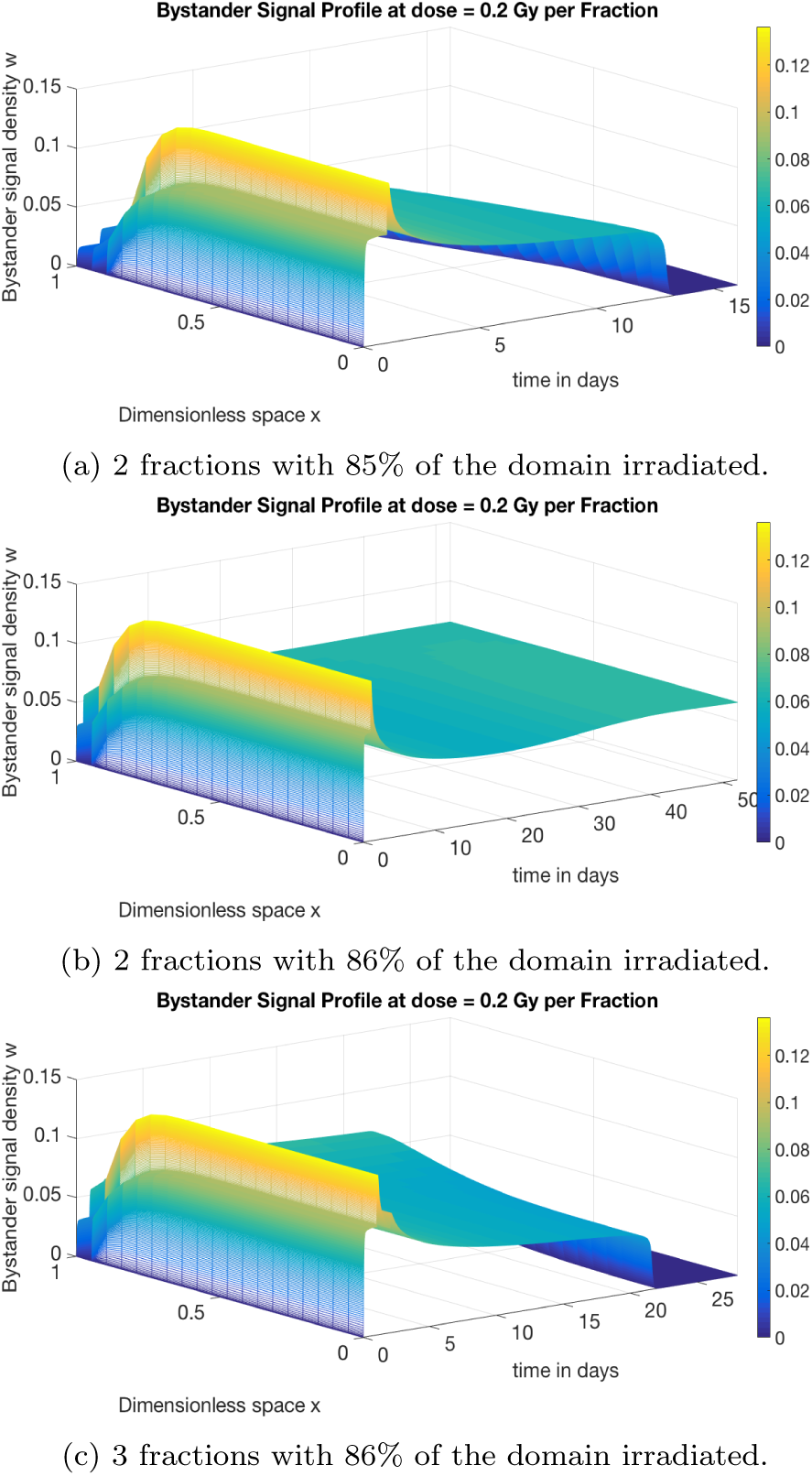
Bystander signal profiles with fractionated radiation exposure in a heterogeneous domain. All simulations done at *A*_0_ = 0.3.

In Figures 10b and 10c we observe the effect of number of exposures on the dynamics of the signal. Two exposures lead to a convergence to a nonzero steady state while three exposures ensure a complete decay of the signal.

There is the need for further understanding of the effects of the number of fractions and the size of irradiated domain on the qualitative behaviors of the bystander signal, and it is time for a phase plane analysis.

## 4 Analysis of the Signal’s Lifespan in a Homogeneous Domain

In this section, we will analyze the model to further understand both the zero and the nonzero steady state convergence observed in the previous section as well as the effect of fractional radiation treatment schedules on the signal’s dynamics. Since the focus is on a homogeneous domain, we ignore the diffusion terms so that the system of PDEs reduces to a system of ODEs. For the moment, we will focus on a single radiation exposure. Since we are interested in the large-time dynamics of the model, we may view all the radiation terms as a one-time input into the model which can be accommodated into the initial conditions. Then system (12) becomes

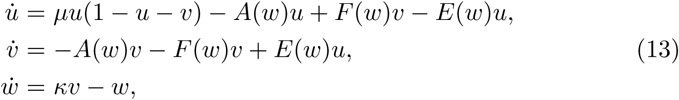

coupled with appropriate initial conditions for *u*(*t*), *v*(*t*), and *w*(*t*), respectively.

We will constrain the initial conditions for the cells to be below their carrying capacity, i.e., *u*(0) + *v*(0) ≤ 1 and consider a very small concentration of bystander signal initially present, i.e., *w*(0) ≪ *𝛋*. These conditions are biologically reasonable for healthy cells because they can not grow beyond their carrying capacity. Under these conditions, the system (13) has the following properties:

### Theorem 1 (Positivity and Boundedness)

*All solutions of the system (13) are positive for all time*, *t*. Moreover, if *u*_0_ = *u*(0) ≥ 0, *v*_0_ = *v*(0) ≥ 0 *and w*0 = *w*(0) ≥ 0, *then*

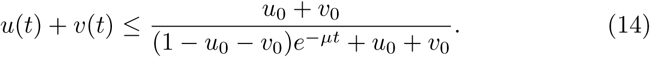

*Furthermore, if u*_0_ + *v*_0_ ≤ 1 *and w*_0_ ≤ κ, *then*

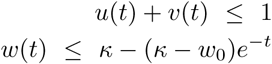

*and w*(*t*) ≤ *κ*, *for all time, t.*

*Proof* Since

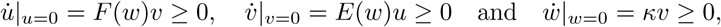

then all solutions are nonnegative for all time, *t*.

Let *m* = *u* + *v* and *m*_0_ = *u*_0_ + *v*_0_. We have

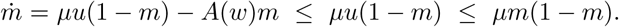

Thus,

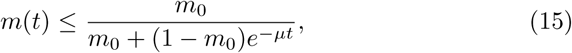

which implies (14) and *u* + *v* ≤ 1 if *u*_0_ + *v*_0_ ≤ 1.

Also, in case of *v* ≤ 1,

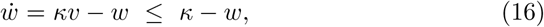

which implies that

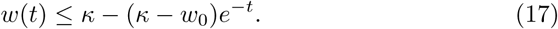

### Corollary 1 (Forward Invariant Region)

*The set*

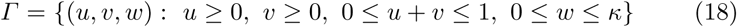

*defines a forward invariant region of the system (13).*

The forward invariant region is closed and bounded in **R**^3^; and therefore compact. If we restrict the phase plane to this invariant region, then any trajectory with initial condition in this region will remain in the region for all times. This suggests that the system of three ODEs (13) has a global attractor.

We can further reduce this system from three ODEs to two ODEs by simply assuming that the bystander signal’s dynamics is faster than cells’ dynamics (a fact that was suggested by the data fitting of the model). This implies that *ẇ* = 0 and we have

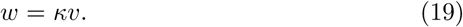

So

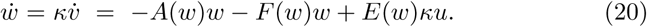

Combining eqs. (13),(19), and (20) we derive

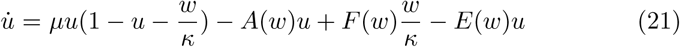

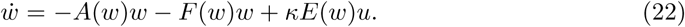

### 4.1 Phase Plane Analysis of the System of Two ODEs

The boundary steady states of (21)–(22) are (0, 0) and (1,0). Interior steady states,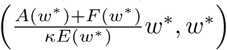 exist provided Ψ(*w*^*^) = 0 and

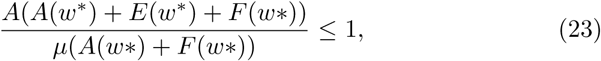

with

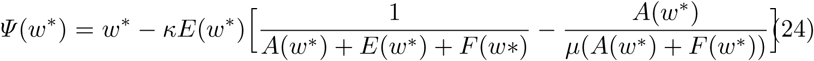

A typical form of *Ψ*(*w*^*^) is shown in Figure 11. We observe that (24) has up to three zeros (including *w*^*^ = 0 which is already listed above) which implies that the system admits between two and four steady states depending on the values of model parameters.

The trivial equilibrium, (0,0), is a saddle, and the homogeneous equilibrium, (1, 0), is a stable node. An interior equilibrium, (*u*^*^, *w*^*^), is stable if

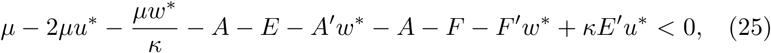

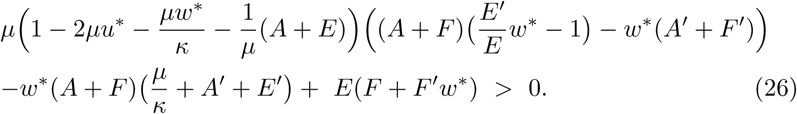

**Fig. 11:**
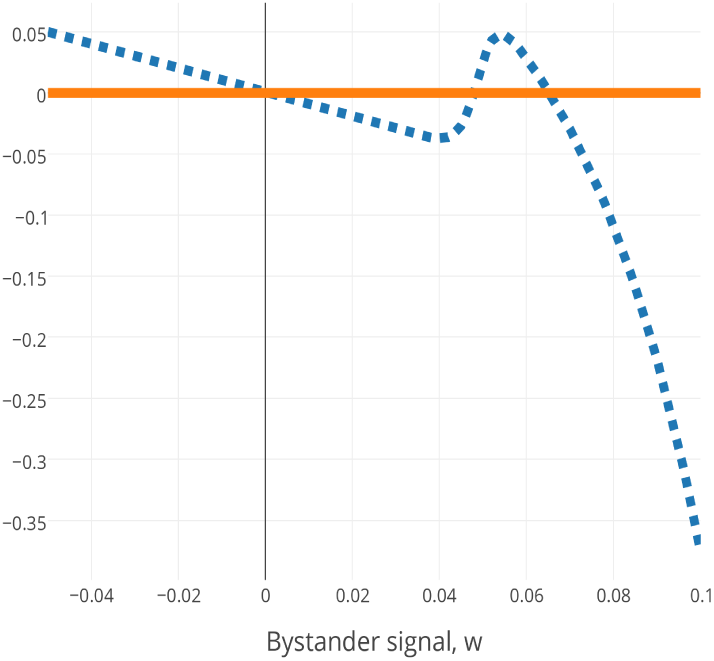
The graph of Ψ(*w*^*^) as defined in (24) for *A*_0_ = 0.3.

Otherwise, (*u*^*^, *w*^*^) is unstable.

The nullclines of the system (21)– (22) are shown in Figure 12 for three different values of the parameter *A*_0_. For *A*_0_ = 0.3, the system has two interior equilibria, one of which is a saddle and the other of which is a stable node, as shown in Fig. 12a. The stable manifold of the interior saddle equilibrium forms a separatrix (dotted line) that demarcates the basins of attraction of the stable interior steady state and the stable homogeneous steady state, (1, 0), into Region I and Region II, respectively. The interior equilibria coalesce and disappear through a saddle-node bifurcation when the condition in (23) is violated at *A*^*^ ≃ 0.58 as shown in Figure 12b and 12c. For *A*_0_ > *A*\*, there are no interior steady state. The homogeneous steady state, (1,0), is the global attractor in this case. Furthermore, we observe that a trajectory that originates from Region I can be pushed into Region II by a further radiation exposure as seen in Fig. 12d. This explains the effect of multiple radiation exposures previously observed in the qualitative behavior of bystander signal profile.

**Fig. 12:**
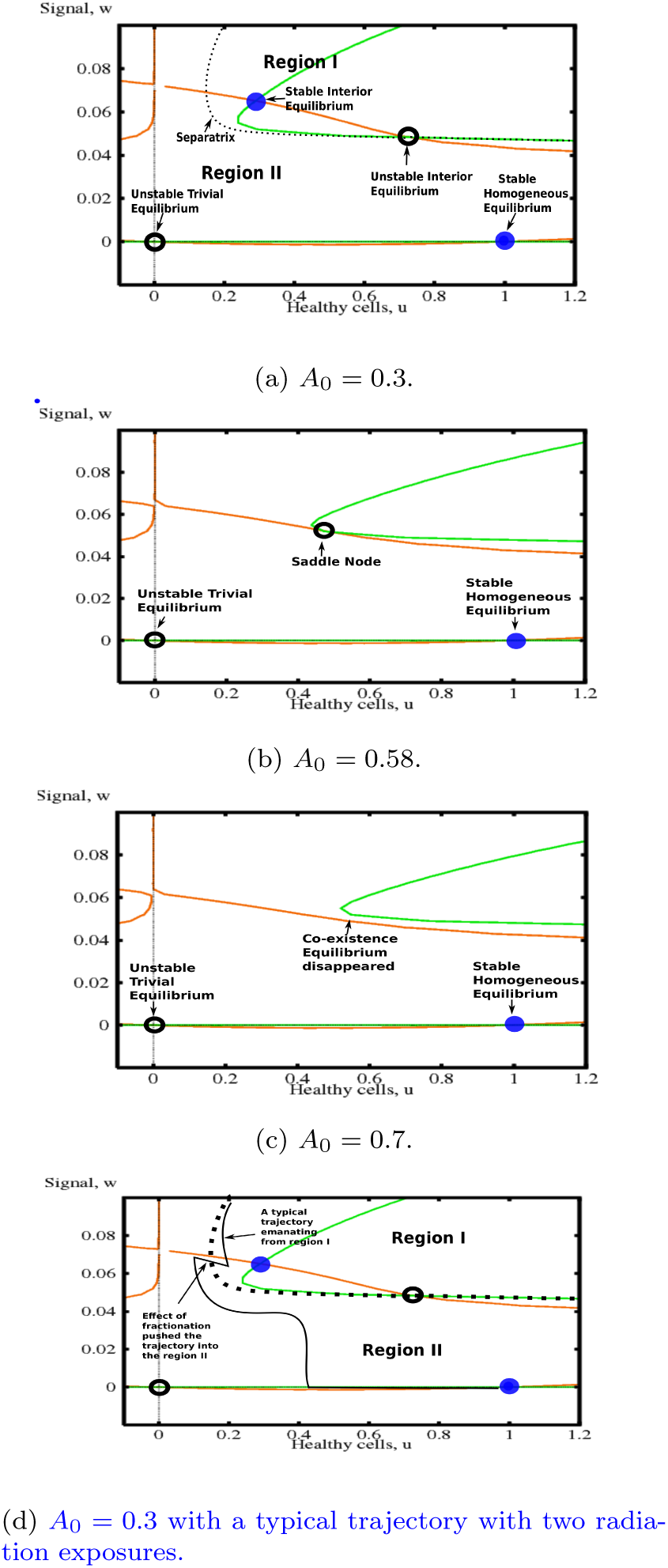
Phase plane showing the nullclines of the system (21)–(22) for three different values of *A*_0_. (a) *A_0_ =* 0.3; (b) *A_0_ =* 0.58; (c) *A_0_ =* 0.7; (d) *A_0_ =* 0.3 with a typical trajectory with two radiation exposures.

In what follows, we want to make connections between the previous phase plane analysis and the *lifespan* of a bystander signal - which we make more precise in the following.

### Definition 1 (Lifespan)

Let *f*_1_, *a*_1_, and *e*_1_ be the lower thresholds of the bystander effects as in Equations (5) – (8). Let *k* = min{*f*_1_, *a*_1_, *e*_1_}.

1. A bystander signal *w*(*t*), at time, *t*, is called active (or inactive) if *w*(*t*) > *k* (or *w*(*t*) < *k*).
2. Suppose that *T* is the time at which the bystander signal becomes active, i.e., *w*(*T*) = *k* and *w*(*t*) < *k* for *t* < *T*. Let

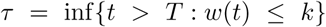

be the time at which the signal, *w*(*t*), first becomes inactive after time, *T*. If *τ* exists, then we define the lifespan of the emitted bystander signal as *τ*. Otherwise, the lifespan is defined by the host’s lifespan.
3. A bystander signal *w*(*t*) is called a transient-state signal if the bystander signal lifespan is finite. Otherwise, it is called a steady-state signal.

Indeed, any trajectory with initial condition in Region I of Fig. 12a converges to the stable interior steady state, while any trajectory with an initial condition in Region II of Fig. 12a persists for a while but eventually converges to zero. We can see this clearly illustrated in Fig. 13a. This persistence is due to the slow transit along the saddle interior equilibrium, which may take many days. However, as the values of *A*_0_ increases and the homogeneous steady state, (1,0), becomes the global attractor, we observe that it takes shorter time for trajectories to converge to the homogeneous steady state, (1, 0). This is illustrated in both Figures 13b and 13c.

**Fig. 13:**
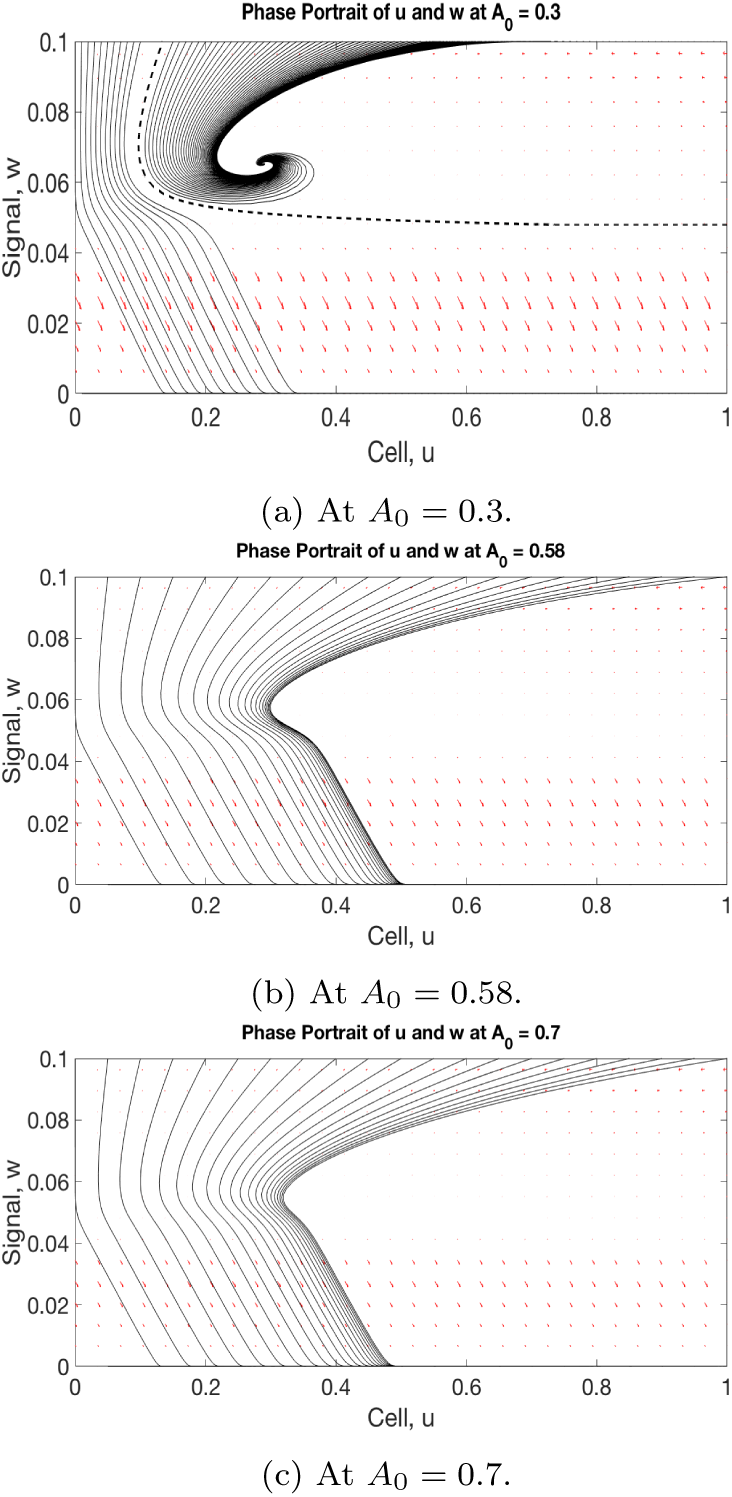
Phase portrait at different values of *A*_0_. *A*_0_ = 0.3; (b) *A*_0_ = 0.58; (c) *A*_0_ = 0.7.

### 4.2 Sensitivity Analysis of the Lifespan of the Bystander Signal

In this section, we identify parameters that affect the lifespan of the bystander signal the most. We study the normalized sensitivity coefficient *S_τ_* of the bystander signal’s lifespan, τ, to a parameter *p*, that is,

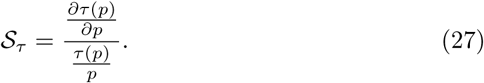

*S_T_* can be interpreted as the % change in the signal’s lifespan per 1% change in the value of a cell’s parameter. *S_τ_* can be positive or negative indicating the parameter increase or decrease the lifespan of the signal.

Since we do not have an explicit formula for the signal’s lifespan, we estimate 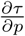 using the central difference approximation:

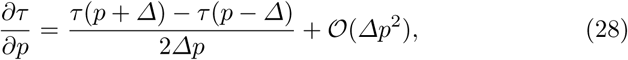

where Δρ = 1% of *p*. The resulting sensitivity indices of the lifespan are summarized in Table 4.

**Table 4:**
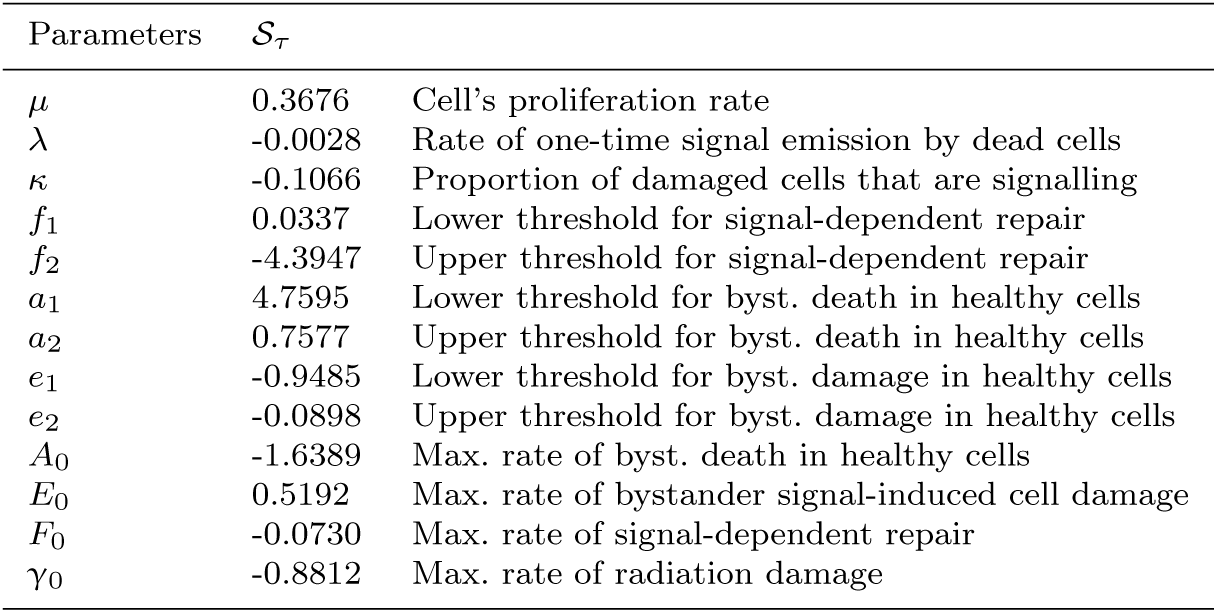
Sensitivity analysis of the bystander signal’s lifespan.

| Parameters | $\mathcal{S}_\tau$ | |
| --- | --- | --- |
| $\mu$ | 0.3676 | Cell's proliferation rate |
| $\lambda$ | -0.0028 | Rate of one-time signal emission by dead cells |
| $\kappa$ | -0.1066 | Proportion of damaged cells that are signalling |
| $f_1$ | 0.0337 | Lower threshold for signal-dependent repair |
| $f_2$ | -4.3947 | Upper threshold for signal-dependent repair |
| $a_1$ | 4.7595 | Lower threshold for byst. death in healthy cells |
| $a_2$ | 0.7577 | Upper threshold for byst. death in healthy cells |
| $e_1$ | -0.9485 | Lower threshold for byst. damage in healthy cells |
| $e_2$ | -0.0898 | Upper threshold for byst. damage in healthy cells |
| $A_0$ | -1.6389 | Max. rate of byst. death in healthy cells |
| $E_0$ | 0.5192 | Max. rate of bystander signal-induced cell damage |
| $F_0$ | -0.0730 | Max. rate of signal-dependent repair |
| $\gamma_0$ | -0.8812 | Max. rate of radiation damage |

The lower threshold for the bystander-induced death in healthy cells, *α*_1_, has the strongest positive relationship to the lifespan of the bystander signal. The positive value suggests that a cell that is highly resistant to signal-induced death, i.e., will emit a longer-lived bystander signal. In contrast to the lower threshold of the signal-dependent repair, *f*_1_, which has the lowest positive sensitivity index, *α*_1_ will be a more important parameter to control in order to reduce the lifespan of an emitted bystander signal.

The upper threshold for the signal-dependent repair, *f*_2_, has the strongest negative relationship to the lifespan of the signal. This is because any increase in *f*_2_ will allow for more repair in the damaged cells and thereby reducing the number of cells emitting the bystander signal. This parameter also is a good candidate to control in order to reduce the lifespan of an emitted bystander signal.

Parameters such as λ, 𝛋, and *γ*_0_ that enhance the production of bystander signal have negative sensitivity indices. This negative relationship to the signal’s lifespan might seem counterintuitive. Since an increase in either λ, *𝛋* or *γ*_0_ will lead to an emission of more signals. However, increasing the bystander signal production will also increase the rate of bystander signal-induced death in cells. This in turn leads to the death of damaged cells, which are the emitters of bystander signals, and ultimately leads to a reduction in the lifespan of the signals.

The rate of proliferation, *μ*, has a positive relationship to the signal’s lifespan. This increased proliferation will ensure a quick re-population of cells after radiation exposure. This will in turn yield a consistent increase in the population of the damaged cells via bystander signal-induced damage leading to a longer-lived signal.

Parameters such as *e*_1_, *e*_2_ and *F*_0_ whose increase in value reduce the population of the damaged cells have negative sensitivity indices. This is because reduced production of bystander signal will quick decay due to the threshold-dependent nature of bystander effects.

## 5 Sensitivity Analysis of the Bystander Signal-induced Cell Death

Bystander signal-induced death is one of the bystander effects in cells that needs to be further understood. Many biological observations and mathematical models have found this bystander effect to be more pronounced with low doses of radiation [30,35]. Our model exhibit the same behaviour, as illustrated in Figure 14. We found that, although the direct radiation-induced death i.e., the direct cell kill, is monotonically increasing with doses, the bystander cell death is non-monotonic. Indeed, we observe more bystander cell death at lower doses than at higher doses. The non-monotonicity of the bystander cell death reflects in the overall effect of radiation, i.e., the total cell kill, where there are more cell death than expected at low doses. This is the phenomenon of hyper-radiosensitivity and increased radio-resistance described in Section 1.1. We use sensitivity analysis similar to the one used in the previous section to investigate the cell parameters that affect this bystander cell death the most.

**Fig. 14:**
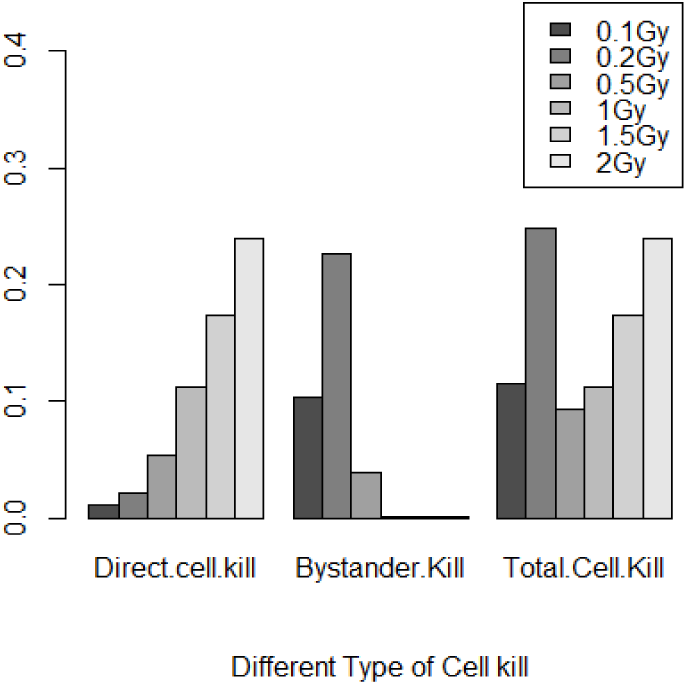
Histogram comparing the total cell death, direct radiation-induced death, and bystander signal-induced death, respectively at different doses.

Let *x*^*^(*t*) be the difference between the cell death at time, *t*, computed using the full bystander model (12) and the cell death at time, *t*, computed when the bystander signal component of the bystander model is removed. The sensitivity coefficient, *S_x_*\*, of the bystander cell death to a parameter, *p*, is given by

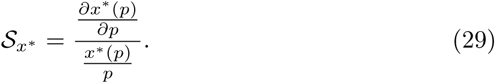

We will use forward difference to estimate 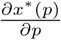. The resulting sensitivity indices of the bystander signal-induced cell death are summarized in Table 5.

**Table 5:** Sensitivity analysis of the bystander signal-induced cell death.

| Parameters | $\mathcal{S}_{x^*}$ | |
| --- | --- | --- |
| $\mu$ | -0.0366 | Cell's proliferation rate |
| $\lambda$ | 0.0044 | Rate of one-time signal emission by dead cells |
| $\kappa$ | 3.4630 | Proportion of damaged cells that are signalling |
| $f_1$ | -0.1792 | Lower threshold for signal-dependent repair |
| $f_2$ | -0.4733 | Upper threshold for signal-dependent repair |
| $a_1$ | -0.9903 | Lower threshold for byst. death in healthy cells |
| $a_2$ | -0.9699 | Upper threshold for byst. death in healthy cells |
| $e_1$ | -0.3306 | Lower threshold for byst. damage in healthy cells |
| $e_2$ | -0.4946 | Upper threshold for byst. damage in healthy cells |
| $A_0$ | 0.4012 | Max. rate of byst. death in healthy cells |
| $E_0$ | 0.2220 | Max. rate of bystander signal-induced cell damage |
| $F_0$ | -0.5066 | Max. rate of signal-dependent repair |
| $\gamma_0$ | 2.7417 | Max. rate of radiation damage |

The parameters such as *𝛋*, *E*_0_ and *γ*_0_ that enhance the population of the damaged cells all have strong relationships to the bystander signal-induced cell death. This is because increase in bystander signal production will in turn increase the rate of bystander signal-induced death in cells. However, *𝛋* has the strongest positive relationship to the bystander signal-induced cell death and will be a more important parameter to control in order to reduce the signal-induced death in cells.

As expected, the signal-induced death in cells increases with increase in the rate of bystander signal-induced death in cells, *A*_0_, as seen in the positivity of its sensitivity index. On the other hand, its lower and upper thresholds, *a*_1_ and *a*_2_, both have negative relationship to the signal-induced death. This is because increase in any of these thresholds will reduce the rate of signal-induced death in cells, which explains their negative sensitivity indices.

Parameters such as *f*_1_, *f*_2_, *e*_1_, *e*_2_ and *F*_0_ whose increase in value reduces the population of the damaged cells all have negative relationship to the signal-induced death in cells. This is simply because fewer damaged cells will result in lower concentration of emitted signals and in turn result in less bystander effects including the signal-induced cell death.

Lastly, the rate of cell proliferation, *μ*, has a negative relationship to the signal-induced death in cells. This seems counterintuitive but increase in cell death will reduce the cell’s population below their carrying capacity. This population reduction will result in cell repopulation which is even more rapid with cells of high proliferation capacity. This repopulation eventually masks the effect of radiation on cells, especially the secondary radiation effects on cell death.

## 6 6 Discussion

Radiation-induced bystander signal is a low-dose phenomenon whose effect on cancer radiotherapy and radiation risk can no longer be overlooked. The dynamics presented in this manuscript have many biological implications.

It is interesting to see that the bystander effect model is able to fully explain the observed HRS and IRR phenomena as seen in the fit in Fig. 6. This is in perfect agreement with the result of a computational model of Powathil et. al. in [35] and experimental observations of Mothersill and Seymour [31]. Although there are other possible explanations of the HRS/IRR phenomena as well, namely, the cell cycle arrest via the G2-checkpoint [27] and the ATM-independent p53-dependent apoptosis [46]. G2 checkpoint is a regulatory mechanism that prevents damaged G2-phase cells from proceeding to mitosis until the damage is repaired. At low doses, this checkpoint is not quickly activated; and damaged cells entering mitosis without repair eventually undergo apoptosis. This leads to an increased cell death at low doses; and hence the phenomenon of HRS/IRR. ATM-independent p53-dependent apoptosis is the apoptotic pathway that is initiated by the tumor suppressor protein, p53, when a damage is irreparable. This mechanism is similar to the mechanism of Cytochrome Complex, except that bystander signal can also involve molecules or proteins which are not p53-dependent. There has been experimental evidence for these three hypotheses, but it is still not clear which, or in which combination, they contribute to the HRS and IRR; and thus further analysis is required. However, our model shows that bystander effect is an important contributor to the HRS/IRR phenomenon.

The parameter estimation in Section 2 computes a 95% confidence interval for each parameter value. The confidence intervals for the lower thresholds of bystander effects considered in this work (that is, bystander signal-induced cell death, DNA repair delay and bystander signal-induced cell damage) present a possible succession of occurrence of these bystander effects. Proper ordering of the lower bounds of these confidence intervals i.e., *f*_1_ ≤ *b*_1_ ≤ *e*_1_ ≤ *α*_1_, gives some insight into the possibility of damaged cells experiencing DNA repair delay as the first bystander effect. This is followed by the bystander signal-induced death in the damaged cells and, afterwards, the bystander signal-induced damage and death in the healthy cells population. This sequence suggests that the damaged cells respond to the signals before the healthy cells respond.

The question about the lifespan of a bystander signal once it is emitted has been questioned in the literature [6]. Mothersill and Seymour in [30] found that an emitted bystander signal can still cause bystander effects in cells even 60h after its emission. Also, Goyanes-Villaescusa [15] and Pant and Kamada [34] observed that the signal can persist for several months and years, respectively. In our model, we found two interior steady states whose stability are respectively stable and saddle. We also found a stable homogeneous steady state at (*u* = 1, *w* = 0). The stable manifold of the saddle interior steady state forms a separatrix that demarcates the basins of attraction of the two stable steady states. We observe that the trajectories that converge to the stable interior steady state provides an explanation for the observation of the presence of bystander signal in cells exposed to radiation several years before. On the other hand, trajectories that converge to the stable homogeneous steady state, (1,0), pass by the interior saddle point leading to a long persistence.

We found a condition for the existence of the interior steady states in Eqn. (11). Although this condition depends on all the bystander effects considered in this work, we also found a dependence on the cell’s proliferation capacity, *μ*. We observe that cells with high values of *μ* are likely to admit the interior steady states while lower values of *μ* are unlikely to satisfy Eqn. (11). Also, we observe the dependence of the existence of this interior steady states on the values of the bystander effects. In particular, we found the maximum rate, *A*_0_, of bystander signal-induced death as an important bifurcation parameter leading to a saddle node bifurcation. Higher values of *A*_0_ leads to disappearance of the interior steady state while the system admits these interior steady states at lower values.

In the absence of the interior steady state, we found several sensitive parameters that can influence the signal’s lifespan. The strongest of these is the lower threshold, α_1_, for bystander signal-induced death in cells. This parameter will be very important in controlling the effect of bystander signal. Most importantly, we also observe that highly proliferating cells emit a longer-lived bystander signal. This is also corroborated by the dependence of the condition in Eqn. (11) on *μ*.

The understanding of the signal’s behavior in a homogeneous environment with multiple radiation exposures is of great interest since fractionated radiation treatment is a very common treatment strategy for cancer. Our model shows that the bystander signal is being produced immediately after each fraction as seen in Section 3. Although the peak of signal concentration does not grow with increase in radiation exposures, the peak of the signal concentration immediately after the second radiation exposure is significantly higher than the rest. We also observe that an increase in radiation exposure can push a trajectory out of the basin of attraction for the stable interior steady state to the basin of attraction of the stable homogeneous steady state. Thus, a further radiation exposure can completely change the behavior of an emitted bystander signal.

Bystander signal dynamics when the domain is partially irradiated is very crucial. This is because tumor cells undergoing radiotherapy are usually surrounded by normal tissues. Under singular and multiple exposures, there seem to be a maximum domain size that needs to be irradiated in order for the signal to die out. We will refer to this as *the maximum domain problem.* Similar to the dynamics in a homogeneous domain, we also observe that further radiation exposure can push a trajectory heading for the stable interior steady state to the basin of attraction of the stable homogeneous steady state. Our simulations reveal that the signal dynamics with more than two fractions will always converge to a zero steady state.

The understanding of the dynamics of the bystander signal cell kill is very interesting to the radiologists in administering low dose of radiation to surrounding normal tissues while the target volume is treated to a high radiation doses. This is usually done in order to reduce radiation toxicity to the surrounding tissues. The sensitivity analysis of the bystander signal cell kill in Section 5 shows that the maximum rate, *E*_0_, of radiation damage, the fraction, *𝛋*, of damaged cells and the maximum rate, *γ*_0_, of radiation damage are very strongly influential on the bystander signal-induced cell death. Thus, highly resistant cells to radiation damage will also be resistant to the bystander cell kill.

In this paper, we analyzed some spatial aspects of the model, but additional analysis certainly would be fruitful. In typical radiation treatments, the radiation dose is not applied uniformly over the tumor domain, hence a distinct spatial structure is imprinted. Also, the bystander signal might be transported differently in different types of tissue such as tumor tissue, stroma, blood vessels, collagen networks etc. However, a detailed consideration of such spatial aspects is beyond the scope of this paper and needs to be left for future research.

Our model incorporated three bystander effects: cell repair delay, bystander signal-induced death and damage. There is the need to incorporate more bystander effects like genetic instabilities, mutation, etc., for more insight and richer understanding of this phenomenon of bystander effects.

## Footnotes

1 The word “delay” in a biological context often refers to the fact that a process is slower than normal. It does not necessarily refer to a delay term as would arise in a delay equation.

## Acknowledgements

We are particularly grateful to G. Powathil for inspiration and discussions about bystander effects, and we thank an anonymous referee for helpful suggestions. We are grateful to helpful suggestions of W. Roda on the implementation of the Goodman and Weare Affine invariant ensemble MCMC sampler for data fitting. We are also grateful for the feedback from the Journal Club of the Center for Mathematical Biology at the University of Alberta. OO is supported by the President’s International Doctoral Award, GdeV and TH are supported by NSERC.

